# Perturbation of the mitochondrial import machinery by aggregation prone Tau affects organelle morphology and reduces neuronal complexity

**DOI:** 10.1101/2022.11.30.518502

**Authors:** Hope I Needs, Kevin A Wilkinson, Jeremy M Henley, Ian Collinson

## Abstract

Protein import into mitochondria is an intricate and highly conserved process essential for organellar biogenesis, and maintenance of its structure and function. Defects in the import apparatus impact the assembly of the respiratory chain and ATP synthase complexes required for oxidative phosphorylation, compromising the supply of ATP to the cytosol. The consequences of reduced bioenergetic function are particularly severe for cells with high energetic demands, such as neurons. However, relatively little is known about how defective import contributes to neurodegeneration, or how aggregation prone toxic proteins, characteristic of neurodegenerative disease, impact mitochondrial import efficiency. Here, we used HeLa cells to investigate how expressing Tau, or a disease-causing variant, affects mitochondrial import activity, morphology, and function. We found that a variant associated with frontotemporal dementia (Tau^P301L^), but not the native version, colocalises with mitochondria, associating with TOM40–the protein-channel component of the outer membrane import complex. Interestingly, Tau^P301L^ production had no discernible effect on overall mitochondrial import function, despite associating with TOM40 and altering mitochondrial morphology. This raised suspicions of a rescue mechanism manifested by the appearance of microtubule and actin containing tunnelling nanotubes (TNTs), used to recruit healthy mitochondria from neighbouring cells and/ or dispose of mitochondria containing aggregated Tau. Furthermore, in primary neuronal cultures Tau^P301L^ induces morphological changes that resemble a neurodegeneration-like phenotype–also mirrored in cells where the import sites are blocked artificially. These results reveal an intriguing link between the production of aggregation prone protein variants, such as Tau^P301L^ and others, with the mitochondrial protein import machinery relevant to neurodegenerative disease.

## Introduction

Mitochondrial protein import is a highly conserved process by which most mitochondrial proteins are transported from where they are synthesised in the cytosol. The process occurs through various pathways ensuring delivery to the desired localisation within the mitochondrion. The translocase of the outer membrane (TOM) complex acts as the major gate for entry into the intermembrane space (IMS). It consists of assembly/stability subunits (TOM5, TOM6, and TOM7), receptor subunits (TOM20, TOM22, and TOM70), and a central pore forming subunit (TOM40) (Chacinska et al., 2009; Tucker & Park, 2019). Proteins traversing through TOM40 are then dispersed *via* distinct pathways, depending on their structure, function, and target destination (Chacinska et al., 2009). The pre-sequence pathway is used by precursor proteins with cleavable N-terminal mitochondrial targeting sequences (MTS). These proteins are either laterally inserted into, or transported across, the inner mitochondrial membrane (IMM); respectively, through the translocase of the inner membrane sort (TIM23^SORT^) or motor (TIM23^MOTOR^) complexes. Cleavage of the MTS enables release and folding of the mature protein (Chacinska et al., 2009).

These pathways (recently reviewed in refs. (Bar-Ziv et al., 2020; Harbauer et al., 2014; Needs et al., 2021)) need to be tightly regulated, and subject to quality control, to maintain mitochondrial structure and function – required for the supply of energy (ATP), cellular homeostasis, and overall health (Sokol et al., 2014). Mitochondrial dysfunction is one hallmark of neurodegeneration, which is being increasingly linked to failure of the protein import machinery (Lin & Beal, 2006; Schon & Przedborski, 2011). For instance, accumulated amyloid precursor protein (APP) within mitochondria from patients with Alzheimer’s disease interacts with the protein-channel components of both outer and inner membranes: TOM40 and TIM23 (Devi et al., 2006). Similarly, mitochondria isolated from the brains of people affected by Huntington’s disease are decorated with the aggregation prone variant of the Huntingtin protein (Htt). Again, Htt associates with the TIM23 complex and was shown to inhibit import as a result (Yano et al., 2014).

Neurofibrillary tangles (NFTs) comprise insoluble aggregations made up primarily of hyperphosphorylated Tau protein. NFTs are characteristic of Alzheimer’s disease, as well as of all other primary and secondary tauopathies (Tapia-Rojas et al., 2019). Tau is highly abundant in neurons and is mostly localised in axons, where it binds to microtubules, promoting their assembly and stability (Binder et al., 1985; Black et al., 1996; Witman et al., 1976). Upon binding to microtubules, Tau modulates the activity of the motor proteins Dynein and Kinesin, which in turn regulates the axonal transport of cargo including mitochondria (Chaudhary et al., 2018; Dixit et al., 2008; Ebneth et al., 1998; Mandelkow et al., 2004; Stamer et al., 2002). However, the ability of Tau to bind microtubules is a highly dynamic process and is dependent on different Tau isoforms, mutations, and post-translational modifications. Due to the vital role of Tau in axonal migration, its aberrant behaviour can bring about neurological pathogenesis (Avila et al., 2004; Barbier et al., 2019; Guo et al., 2017; Szabo et al., 2020). The Tau^P301L^ variant is commonly found in patients with tauopathies and has been well characterised in disease models (Goedert & Jakes, 2005; Poorkaj et al., 2001). Critically, Tau^P301L^ is known to have an increased propensity to form NFTs (Murakami et al., 2006).

Mitochondrial function is impaired by abnormal, pathological Tau, but the mechanism is not yet fully understood. The accumulation of Tau into NFTs can elicit several specific mitochondrial effects. These include increased retrograde transport of mitochondria (*i.e.*, towards the soma); reduced complex I activity, ATP levels, and membrane potential; enhanced oxidative stress; and defective mitophagy (Hu et al., 2016; Lasagna-Reeves et al., 2011; Li et al., 2016; Stamer et al., 2002). Furthermore, various studies have shown that aggregation prone Tau accumulates within the IMS and OMM (Cieri et al., 2018; Hu et al., 2016). Tau mitochondrial accumulation and alterations in membrane potential suggest that there may also be import defects, since membrane potential is required for translocation of proteins across the IMM. However, no studies have yet directly examined import in the presence of pathological Tau.

Here, we show that the disease-linked variant, Tau^P301L^, interacts with the channel forming subunit of the TOM complex (TOM40). Tau^P301L^ association with TOM40 corresponds to changes in mitochondrial morphology together with a reduction in neuronal complexity. Interestingly, these changes to neuronal structure were also induced when mitochondrial import sites were artificially blocked. This suggests that perturbation of mitochondrial protein import, specifically by blockage of the TOM channel by variants of Tau or other aggregation prone proteins, may be an important factor in the onset of neurodegenerative diseases.

## Results

### The aggregation prone variant Tau^P301L^ associates with the mitochondrial import machinery

The association of Tau variants with mitochondria was investigated in HeLa cells conditioned with galactose (HeLaGAL); galactose impairs glycolytic metabolism and drives cells to be more dependent on mitochondria for their energy supplies (Aguer et al., 2011). We expressed Myc-Tau^WT^, Myc-Tau^P301L^, or a GFP control in these cells and carried out Western blotting analysis on the fractionated mitochondria and cytosol (Fig. 1A, left panel). The results show that in Myc-Tau^P301L^ containing cells there was a higher proportion of the Tau protein localised to mitochondria compared to those expressing Myc-Tau^WT^ or GFP (Fig. 1A, left panel – quantified in 1B). To test whether the increased mitochondrial localisation of Tau^P301L^ correlates with altered import pathway components, we expanded this analysis to include a selection of subunits of the import machinery (Fig. 1A, right panel – quantified in 1C). Overall, mitochondria from cells overexpressing Tau^P301L^ had lower levels of mitochondrial TOM20 (receptor subunit of the TOM complex of the OMM) and TIM23 (channel forming subunit of the TIM23 complex in the IMM) compared to control cells, but there was no change in abundance of TOM40 (channel forming subunit of TOM complex of the OMM). Together, these data are indicative of mitochondrial import defects induced by Tau^P301L^.

**Fig. 1.**
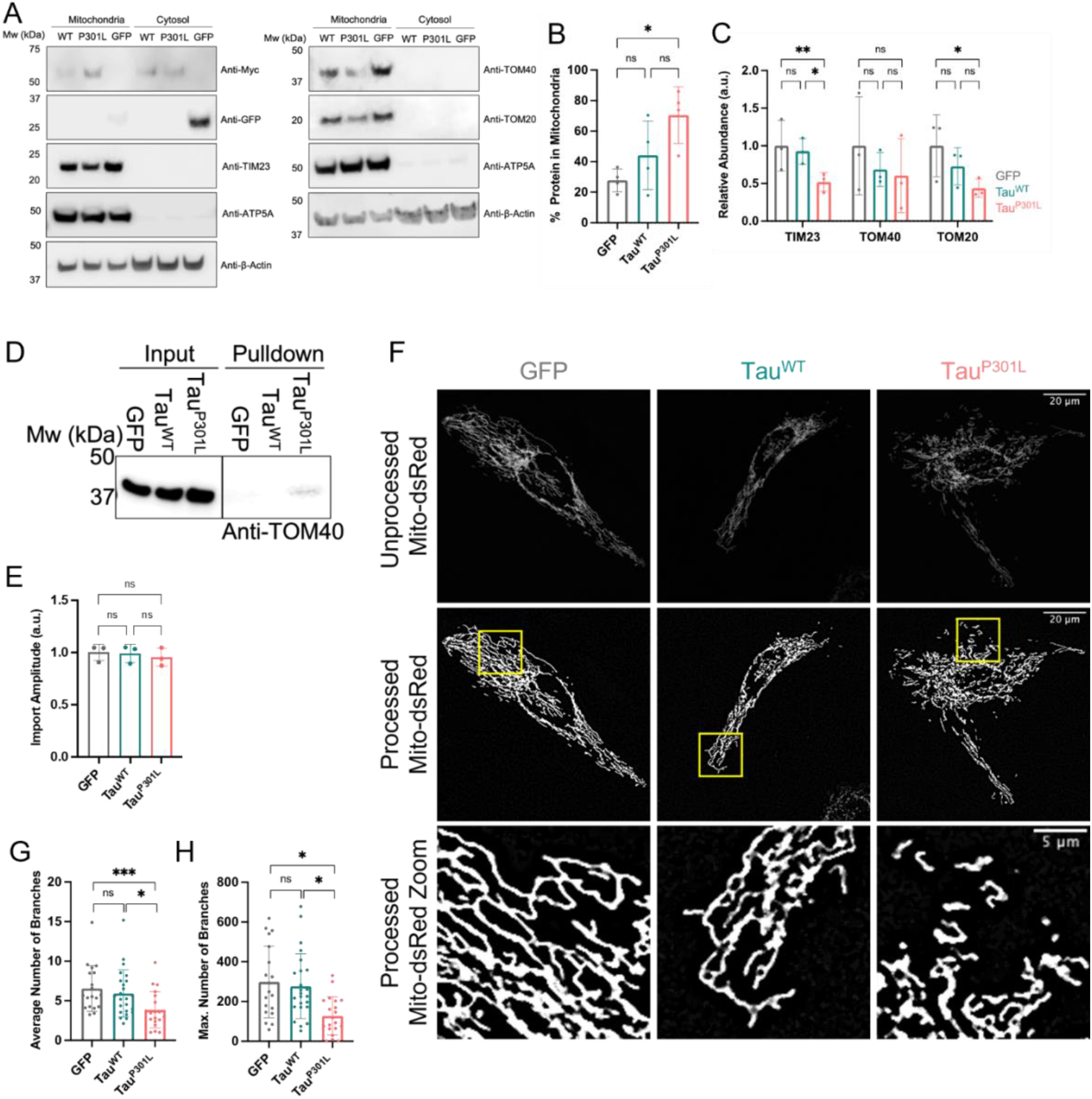
Tau^P301L^ associates with the mitochondrial translocation machinery and alters mitochondrial morphology. **(A)** Representative Western blots showing relative protein abundance in mitochondrial and cytosolic fractions of HeLaGAL cells expressing GFP, Myc-Tau^WT^ or Myc-Tau^P301L^. Localisation of GFP, Tau^WT^ and Tau^P301L^ was analysed by probing for GFP (Sigma; G1544) or Myc (CST; 2276). Translocase subunit abundance was analysed by probing for TIM23 (Invitrogen; PA5-71877), TOM40 (Thermo; PA5-57575), and TOM20 (Santa Cruz Biotechnology; sc-17764). ATP5A (Abcam; 15H4C4) and β-actin (Sigma; A2228) were used as loading controls for mitochondria and cytosol, respectively. N=4 (GFP/Myc) and N=3 (translocases) biological replicates. **(B)** Quantification of the percentage of GFP, Tau^WT^ or Tau^P301L^ in mitochondrial fraction. Normalised to loading controls. Error bars show SD. One-way ANOVA and Tukey’s *post hoc* test were used to determine significance. **(C)** Quantification of translocase subunits (TIM23, TOM40, TOM20). Normalised to loading controls. Error bars show SD. One-way ANOVA and Tukey’s *post hoc* test were used to determine significance. **(D)** Representative Western blot showing TOM40 association with Tau^P301L^, but not with Myc-tagged GFP or Tau^WT^, in mitochondrial fraction of HeLaGAL cells. Mitochondria were isolated from HeLaGAL cells expressing Myc-GFP, Myc-Tau^WT^, or Myc-Tau^P301L^ and proteins lysed with GDN (input), prior to immunoprecipitation using Myc-trap beads (Chromotek). Eluted proteins (pulldown) were analysed by Western blot and probed against TOM40. N=4 biological replicates. Control blot shown in supplementary Fig. S1. **(E)** Maximum import amplitude from NanoLuc import assay on HeLaGAL cells expressing GFP, Tau^WT^, or Tau^P301L^. Bars represent relative maximum mitochondrial import of the precursor protein *Su9-EGFP-pep86* (normalised to eqFP670 expression, max amplitude/run, and control (GFP)). N=3 biological repeats, each with n=3 technical replicates. Error bars display SD. One-way ANOVA with Tukey’s *post hoc* test were used to determine significance. Import traces shown in supplementary Fig. S2. **(F)** Representative confocal images showing mitochondrial morphology in HeLaGAL cells expressing mito-dsRed and GFP (left, grey), Tau^WT^ (middle, teal) or Tau^P301L^ (right, pink). Top panel shows mito-dsRed (mitochondria) prior to processing. Middle panel shows mitochondria after processing, and bottom panel shows a zoom of an area of mitochondria (highlighted in middle panel yellow box) to give a clear view of mitochondrial morphology. N=5 biological repeats. **(G)** Quantification of the average number of mitochondrial branches. Representative images shown in *(F)*. Each point represents an individual cell from a separate z-stack. N=20, 24, 23 for different conditions, respectively, taken from 5 independent biological repeats (N=5). Error bars show SD. Nested one-way ANOVA and Tukey’s *post hoc* test were used to determine significance. **(H)** Quantification of the maximum number of mitochondrial branches in a network. As in *(G).*

Next, we investigated the interaction between Tau^P301L^ and the translocation machinery. We used Myc-trap pull-down assays on mitochondria isolated from cells producing recombinant Myc-GFP, Myc-Tau^WT^, or Myc-Tau^P301L^. Western blots showed that Tau^P301L^, but not its native counterpart or the GFP control, enabled the recovery of detectable quantities of TOM40 (but not TIM23 or TOM20) from the mitochondria of HeLaGAL cells (Fig. 1D; S1). This is consistent with previous reports that showed Tau^P301L^ accumulation in the OMM and IMS (Cieri et al., 2018; Hu et al., 2016), and suggests that it may be associated with the translocation channel in a similar way to an artificially trapped precursor protein (Chacinska et al., 2003; Ford et al., 2022; Hope I Needs et al., 2022). Surprisingly, however, despite association of Tau^P301L^ with the import machinery, in-cell NanoLuc import assays (Hope I. Needs et al., 2022) showed that the import kinetics of a precursor reporter were not significantly affected (Fig. 1E; S2).

Analysis of mitochondrial branching showed that cells expressing Tau^P301L^ have, on average, significantly fewer branches per mitochondrial network when compared to mitochondria from cells expressing Tau^WT^ or GFP (3.9 compared to 5.9 and 6.6, respectively; Fig. 1F, G). The maximum number of branches per network was also reduced to 127.3 for Tau^P301L^, compared to 276.9 for Tau^WT^ and 298.3 for GFP (Fig. 1H). Mitochondrial stress tests measuring oxygen consumption and membrane potential (TMRM fluorescence), showed that despite Tau^P301L^ induced alterations in mitochondrial morphology, respiratory capacity was unaffected (Fig. S3; S4).

### Tau^P301L^ overproduction induces TNT formation

In a parallel study, we explored the effects of artificially blocking the import machinery (Hope I Needs et al., 2022). This can be achieved with the small molecule inhibitor MB20 (Cheung, 2017), or with a precursor fused to DHFR which, when bound to methotrexate, becomes trapped in the channel (Chacinska et al., 2003; Ford et al., 2022). While these treatments inhibit import in isolated mitochondria (Cheung, 2017; Ford et al., 2022), we unexpectedly found that they fail to do so within whole cells (Hope I Needs et al., 2022). We wondered if the apparent lack of effect on the whole mitochondrial population of the cell could be explained by a mitochondrial replacement system. We found that, at the whole cell level, defective mitochondrial function is rescued by the recruitment of healthy mitochondria from surrounding cells, and disposal of compromised mitochondria, by way of TNTs (Hope I Needs et al., 2022). Therefore, we were curious if the association of the Tau variant with the import apparatus induced a similar response.

HeLa cells co-expressing mCherry with either Myc-GFP, Myc-Tau^WT^, or Myc-Tau^P301L^ were analysed by confocal microscopy. The results show that in mitochondrial dependent HeLaGAL cells (conditioned with galactose) production of Myc-Tau^P301L^ induced TNTs: 62% of all cells exhibited TNTs, compared to 22% of cells producing native Tau and 18% of those expressing GFP (Fig. 2A, B). In contrast, glycolytic HeLaGLU cells (cultured in glucose containing media) were significantly less prone to TNT production. These observations could explain why the association of Tau^P301L^ with the import machinery has no impact on protein import activity, due the existence of a TNT-dependent rescue mechanism replacing defective mitochondria in the Tau^P301L^ expressing cells. Consistent with this hypothesis, we observed mitochondria inside the TNTs (Fig. 2C), potentially on their way to rescue Tau affected cells.

**Fig. 2.**
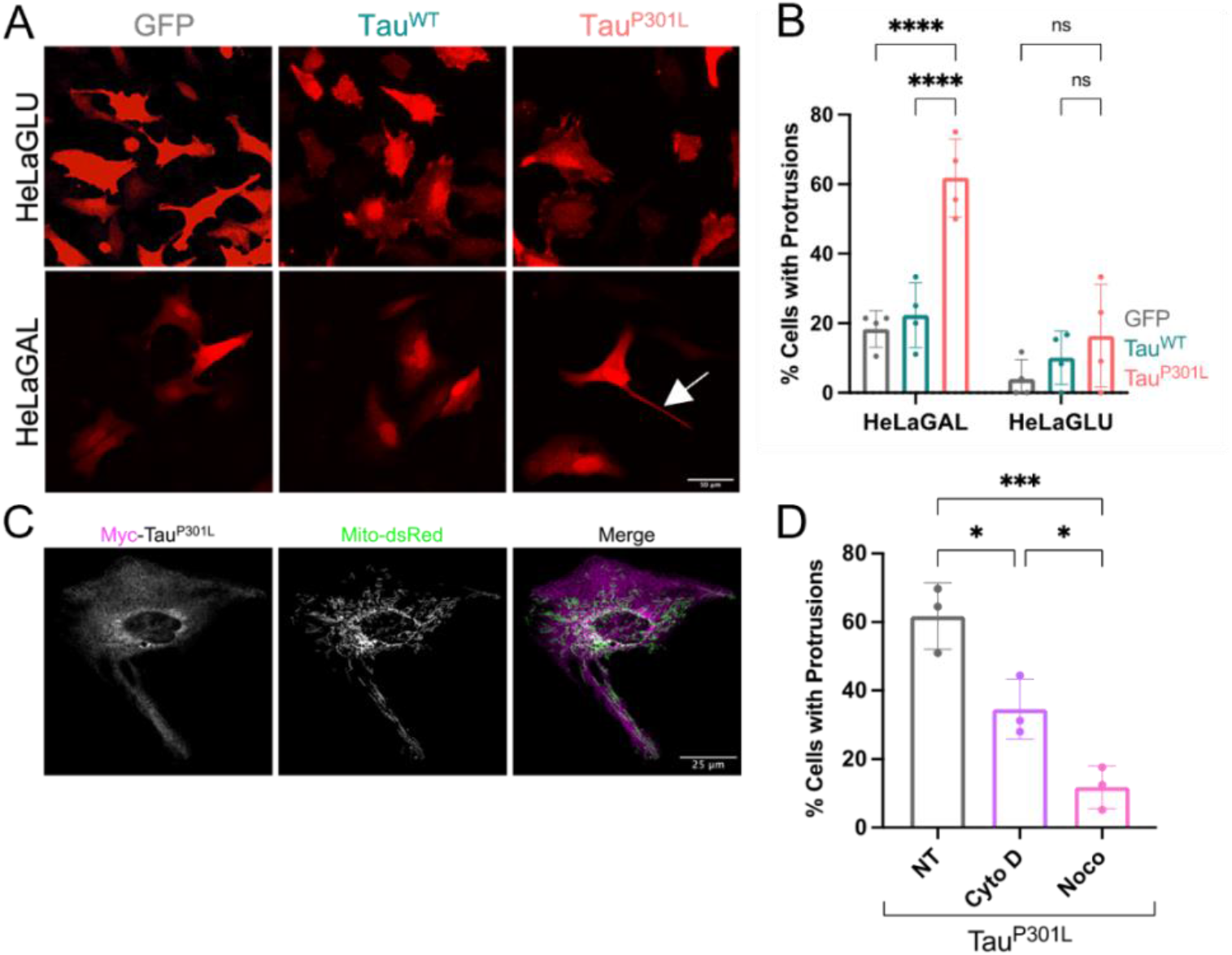
HeLaGAL cells expressing Tau^P301L^ form TNTs that contain mitochondria. **(A)** Representative confocal images showing cell morphology of HeLaGLU or HeLaGAL cells subjected to expression of mCherry (red) as well as GFP, Tau^WT^, or Tau^P301L^. White arrow in bottom right panel indicates a TNT. N=4 biological replicates. **(B)** Quantification of the proportion of cells with protrusions from representative images shown in *(A)*. Error bars show SD. Two-way ANOVA and Tukey’s *post hoc* test were used to determine significance. N=4 biological replicates, 20 cells counted per replicate. **(C)** Representative confocal images showing mitochondrial localisation (mito-dsRed; green) in cells overexpressing Myc-Tau^P301L^ (magenta). N=4 biological replicates. **(D)** Quantification of the percentage of HeLaGAL cells with protrusions following Tau^P301L^ overexpression in the presence of TNT inhibitors. HeLaGAL cells were subjected to expression of Tau^P301L^-mScarlet and either untreated (NT; DMSO only) or treated with 50 nM Cytochalasin D or 100 nM Nocodazole for 48 h. N=3 biological replicates, 20 cells counted per replicate. Error bars show SD. One-way ANOVA and Tukey’s *post hoc* test were used to determine significance. Representative images shown in supplementary Fig. S5.

TNTs induced by import impairment contain actin as well as microtubules, which are necessary for mitochondrial movement (Hope I Needs et al., 2022). Thus, drugs that target actin (cytochalasin D) and tubulin (nocodazole) polymerisation inhibit TNT formation (Bittins & Wang, 2017; Kumar et al., 2017; Luchetti et al., 2012). The TNTs induced here also contain actin and tubulin and their drug response is the same (Fig. 2D; S5). Therefore, perturbation of the import machinery by precursor trapping, small molecule inhibition (shown previously (Hope I Needs et al., 2022)) and, as we show here, aggregation of Tau all seem to induce a common response involving TNTs.

### The overproduction of Tau^P301L^ and precursor trapping both bring about reduced neuronal complexity

Next, we explored the effects of perturbing the mitochondrial import machinery in neurons. In particular, we wanted to compare the effects of precursor trapping (deploying a precursor-DHFR fusion in the presence of MTX) against production of Tau^P301L^. We used confocal microscopy to assess morphology of DIV21 (21 days *in vitro*) primary hippocampal neurons following 7 days of precursor trapping or Tau variant over-production. DIV21 neurons equate to ‘mature’ neurons with spines and fully formed synapses (Grabrucker et al., 2009; LaBarbera et al., 2021) and therefore allow analysis of synapses and neuronal complexity. Axons were identified by staining for Ankyrin G, a scaffolding protein important in formation of the initial segment of the axon (Grubb & Burrone, 2010).

Over-production of native Tau caused a significant reduction in the number of processes per cell compared to the GFP control, while expression of the aggregation prone Tau^P301L^ caused a more pronounced reduction in processes (Fig. 3A, B). Interestingly, despite the reduction in neuronal complexity, neither Tau variant affected axonal length (Fig. 3C). To determine if these effects were attributable to perturbation of the import machinery by Tau, we also subjected primary neurons to mitochondrial precursor trapping. Inclusion of the precursor alone did not affect cell complexity, but when the import of this precursor was stalled by MTX addition, the number of processes was significantly reduced (Fig. 3D, E), mirroring the effect of Tau. In addition, however, stalling import significantly reduced axonal length (Fig. 3F; +MTS, +MTX), suggesting a more severe response. Importantly, the changes in length and complexity were not due to non-specific effects of MTX, as they are dependent on the MTS (Fig. 3E, F; -MTS, ±MTX). Neuronal viability was unaffected by precursor trapping (±MTS) or production of Tau variants, although there was a measurable effect of the methotrexate drug, which is inconsequential for this analysis (Fig. S6).

**Fig. 3.**
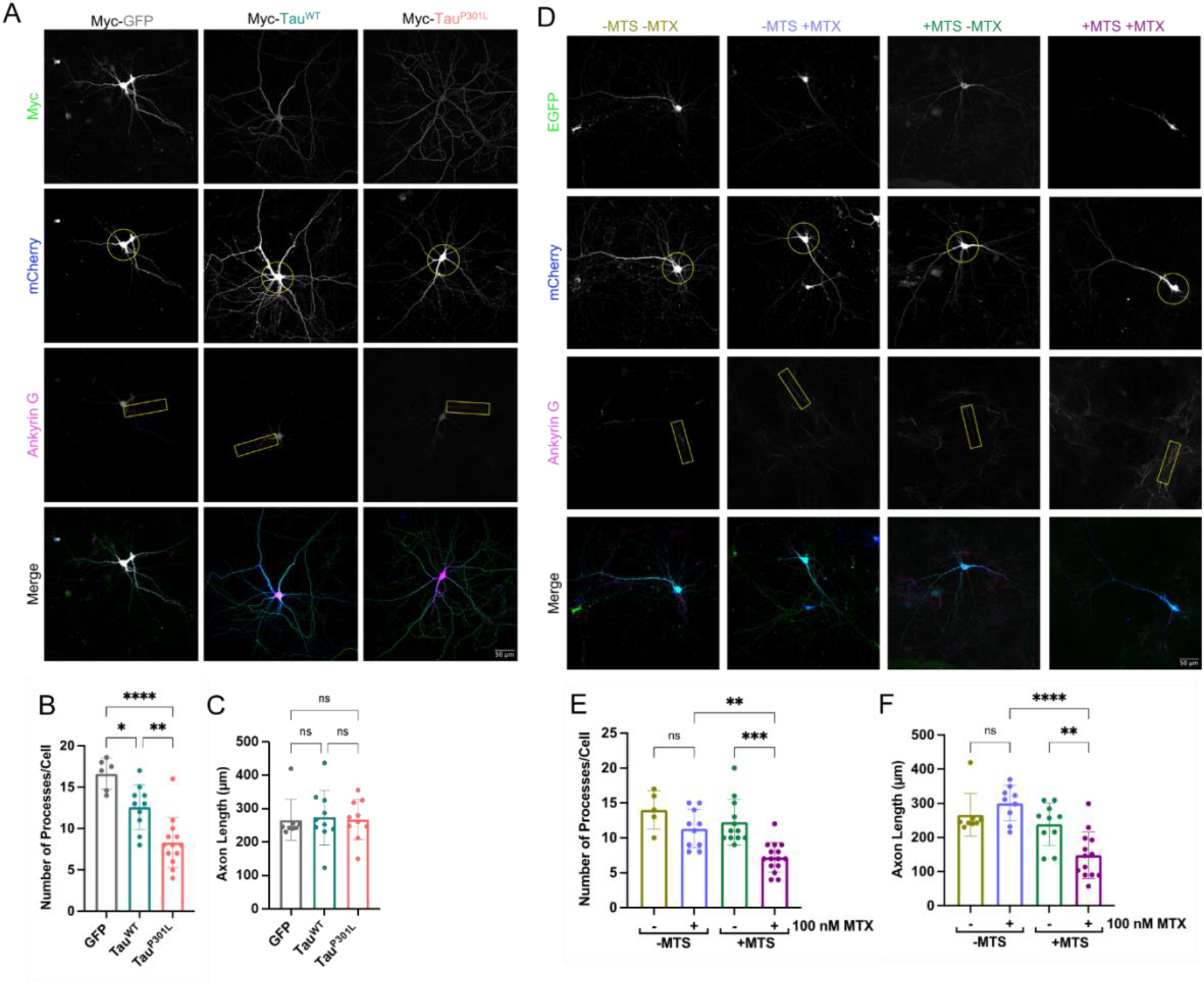
Tau^P301L^ and precursor trapping reduce neuronal complexity. **(A)** Representative images showing morphology of DIV21 primary hippocampal neurons expressing mCherry (whole cell cytosolic marker; blue; circle used to highlight complexity in second panel down *i.e.,* the number of processes that cross the circle is representative of number of processes per cell) and Myc-tagged GFP, Tau^WT^, or Tau^P301L^ (Myc; green), taken by confocal microscopy. Axon initial segments were stained with Ankyrin-G (magenta; highlighted by box in third panel down; NeuroMab 75-146). N=4 biological replicates. **(B)** Quantification of number of processes per cell for GFP, Tau^WT^ and Tau^P301L^ expressing neurons. Each data point represents an individual cell (cells were only analysed if the entire cell could be identified separately from surrounding cells). Error bars show SD. Nested one-way ANOVA and Tukey’s *post hoc* test were used to determine significance. **(C)** Quantification of axon length for GFP, Tau^WT^ and Tau^P301L^ expressing neurons. Cells were only analysed if the axon could be clearly identified. Error bars show SD. Nested one-way ANOVA and Tukey’s *post hoc* test were used to determine significance. **(D)** Representative confocal images showing the morphology of DIV21 primary hippocampal neurons expressing mCherry (whole cell cytosolic marker; blue; circle used to highlight complexity in second panel down *i.e.,* the number of processes that cross the circle is representative of number of processes per cell) as well as EGFP-DHFR (-MTS; green; top panel) or Su9-EGFP-DHFR (+MTS; green; top panel) in the presence or absence of 100 nM MTX. Axons were stained with Ankyrin-G (magenta; highlighted by box in third panel down). N=5 biological replicates. **(E)** Quantification of the number of processes per cell in neurons expressing EGFP-DHFR (-MTS) or Su9-EGFP-DHFR (+MTS) in the presence or absence of 100 nM MTX. Each data point represents an individual cell (cells were only analysed if the entire cell could be identified separately from surrounding cells). Error bars show SD. Nested one-way ANOVA and Tukey’s *post hoc* test were used to determine significance. **(F)** Quantification of axonal length in neurons expressing EGFP-DHFR (-MTS) or Su9-EGFP-DHFR (+MTS) in the presence or absence of 100 nM MTX. Cells were only analysed if the axon could be clearly identified. Error bars show SD. Nested one-way ANOVA and Tukey’s *post hoc* test were used to determine significance.

These results indicate that a reduction in neuronal complexity and length is brought about by perturbation of the mitochondrial protein import machinery. Notwithstanding the differing response to axonal length, this morphological change can be induced either by artificial precursor stalling, or Tau^P301L^ expression, which we assume is mediated *via* a common mechanism.

### Precursor trapping and Tau^P301L^ correlate with fewer synapses

The neuronal analysis was expanded to investigate the number of synapses per dendrite. The major scaffold protein PSD95 is involved in the organisation of excitatory postsynaptic signalling complexes in neurons (Broadhead et al., 2016), and is therefore commonly used as a postsynaptic marker. These signalling complexes comprise glutamate receptors, ion channels, signalling enzymes and adhesion proteins vital in neurotransmission (Chen et al., 2011).

Imaging and PSD95 staining showed that neurons producing Tau^P301L^ contained on average of 1.8 synapses per 10 μm dendrite, significantly lower than cells expressing GFP or native Tau, both of which had about twice as many (Fig. 4A, B). Subsequent Western blotting analysis of whole lysates from cortical cells expressing GFP, or either of the Tau variants, determined that there was no change in the overall abundance of the synaptic markers PSD95, Gephyrin, or Synaptophysin (Fig. S7). This indicates that the noted reduction in synapse abundance, must be due to a change in the distribution of the synaptic assembly protein PSD95 (and possibly others), rather than its quantity.

**Fig. 4.**
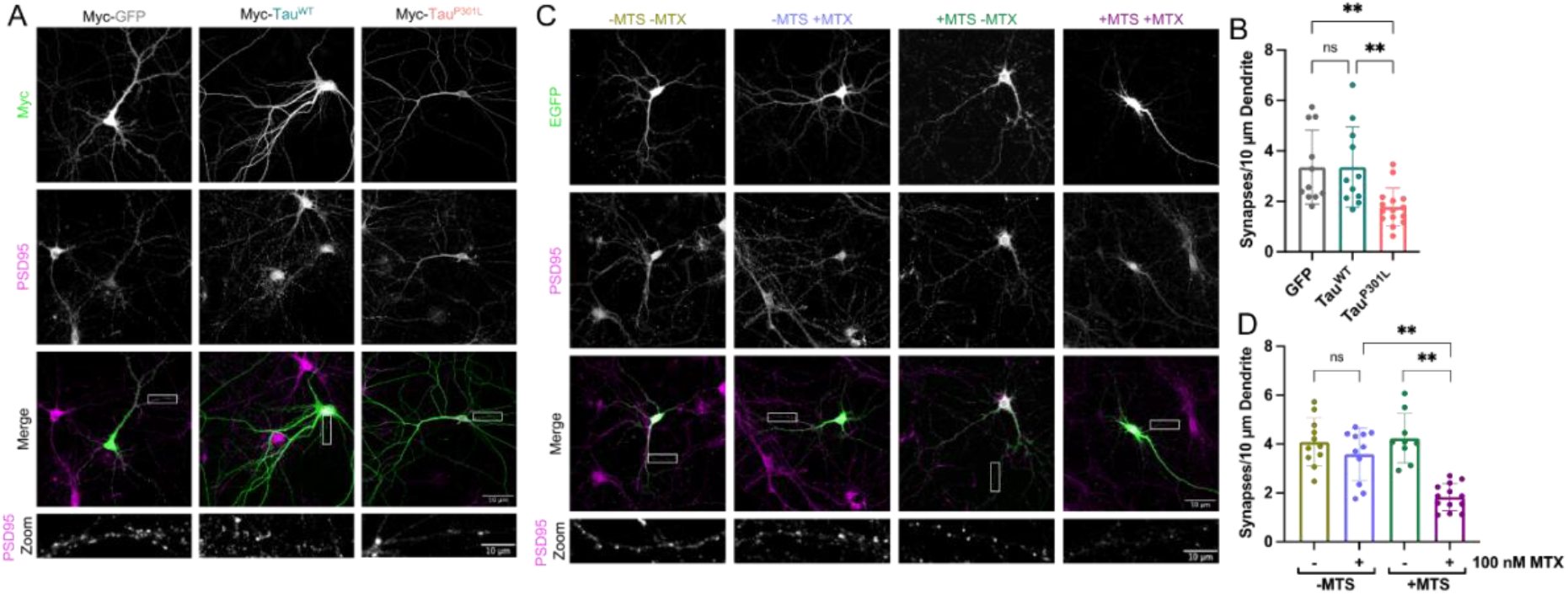
Tau^P301L^ and precursor trapping reduce synapse abundance. **(A)** Representative confocal images showing synaptic staining of DIV21 primary hippocampal neurons expressing Myc-tagged GFP, Tau^WT^, or Tau^P301L^ (Myc; green). PSD95 staining shows synapses (magenta; second panel from top, also shown in zoom; EMD Millipore AB1596). N=5 biological replicates. **(B)** Quantification of the average number of synapses per 10 μm dendrite for GFP, Tau^WT^ and Tau^P301L^ expressing neurons. Synapses were counted manually on blinded data. Each data point represents an individual cell. Error bars show SD. Nested one-way ANOVA and Tukey’s *post hoc* test were used to determine significance. **(C)** Representative confocal microscopy images showing synaptic staining of DIV21 primary hippocampal neurons expressing EGFP-DHFR (-MTS) or Su9-EGFP-DHFR (+MTS; EGFP shown in green, top panels) in the absence or presence of 100 nM MTX (−/+MTX). PSD95 staining shows synapses (magenta, second panel from top, also shown in zoom). N=5 biological replicates. **(D)** Quantification of the average number of synapses per 10 μm dendrite for neurons expressing EGFP-DHFR (-MTS) or Su9-EGFP-DHFR (+MTS) in the presence or absence of 100 nM MTX. Synapses were counted manually on blinded data. Each data point represents an individual cell. Error bars show SD. Nested one-way ANOVA and Tukey’s *post hoc* test were used to determine significance.

Perturbation of the import machinery by precursor trapping induced an almost identical reduction in the number of synapses per dendrite to expression of Tau^P301L^ (Fig. 4C, D). Again, the controls show that synaptic loss is correlated to the precursor trapping and not MTX treatment, since in the absence of a presequence (−MTS) there is no significant change in the number of synapses in the presence of MTX (Fig. 4D, −MTS, ±MTX). Western blotting analysis showed that the overall cellular abundance of all three synaptic marker proteins remained constant irrespective of precursor trapping (Fig. S8), as noted above for the Tau variants. These results demonstrate that expression of aggregation prone Tau^P301L^, which binds specifically to the mitochondrial import machinery, produces defects in neuronal complexity and synapse number similar to those observed upon direct blockade of import.

## Discussion

This study explores aspects of the mitochondrial and cellular responses to Tau, specifically aggregation prone Tau^P301L^ that is implicated in neurodegeneration. We established that association of Tau^P301L^ with the mitochondrial protein import machinery correlates with disease-like phenotypic changes in mitochondria and neurons. Moreover, the effects of Tau mirror those elicited by a precursor protein artificially stalled within the import machinery by a C-terminal fusion of DHFR bound to methotrexate (MTX), suggesting that Tau^P301L^ exerts its effects through perturbation of the mitochondrial import machinery.

At the mitochondrial level, import perturbation by various means cause similarly profound morphological changes including decreased branching (Tau aggregation) and increased fission (precursor trapping) (Hope I Needs et al., 2022). Unexpectedly, however, neither Tau^P301L^ nor precursor trapping affected overall mitochondrial import activity. We postulate that this can be explained by both Tau^P301L^ and precursor trapping inducing the formation of actin and tubulin containing tunnelling nanotubes (TNTs) (Hope I Needs et al., 2022). It has been shown in several previous studies that TNTs can provide a conduit for exchange of mitochondria between cells (summarised in a recent review (Liu et al., 2021)). Thus, we propose that healthy mitochondria are recruited to damaged cells *via* TNTs from nearby unaffected/ healthy cells. Meanwhile, damaged mitochondria are likely disposed of also through TNTs, as we recently showed following precursor trapping (Hope I Needs et al., 2022). This process could also provide a rescue mechanism that explains the observed preservation of mitochondrial import activity and respiratory function despite Tau^P301L^ accumulation within the TOM40 import channel.

The Tau variant induced neuronal changes seen here recapitulate those observed after precursor trapping, strongly suggesting that they occur, at least in part, as a result of perturbation of the import machinery. In this context, we do not yet know if the TNT-dependent rescue response, noted in mitochondria dependent HeLa cells, occurs in neurons. Although we do know that neurons form TNTs, which have been shown to transport Tau itself (Tardivel et al., 2016). Perhaps the rescue mechanism described incorporates, in addition to inter-cellular mitochondrial sorting, routes for the clearance of pathological proteins from critical cells like neurons.

The decrease in processes and synapse number induced by perturbation of the import machinery are consistent with changes observed in neurodegeneration (Coleman & Perry, 2002; Kweon et al., 2017; Wishart et al., 2006). Therefore, an important future goal is to determine if the TNT rescue mechanism is implemented in neurons exposed to import defects, and if so to what extent it mitigates against more severe neuronal aberrations, or cell death. Indeed, the absence of an impact of Tau^P301L^, or precursor trapping, on cell viability may have been due to the rescue process.

The variant Tau^P301L^ has been extensively characterised (Goedert & Jakes, 2005; Murakami et al., 2006; Poorkaj et al., 2001) and, in agreement with the results shown here, it has previously been shown to localise to the mitochondrial IMS and OMM (Cieri et al., 2018; Hu et al., 2016). However, its interaction with the translocation machinery had not previously been reported. Nonetheless, its association with mitochondrial translocases is reminiscent of other proteins implicated in neurodegeneration, such as APP, Huntingtin, and alpha-synuclein (Devi et al., 2006; Devi et al., 2008; Yano et al., 2014). Therefore, the neuronal effects of import perturbation shown here point to a common mechanism that could be an important underlying factor in multiple neurological diseases. Indeed, recent reports have linked failed import or aggregation of proteins within import sites with neurodegenerative diseases: both aggregation prone APP and a disease-associated Huntingtin protein variant become trapped in import sites and are directly correlated with mitochondrial dysfunction, a characteristic feature of neurodegeneration (Devi et al., 2006; Yano et al., 2014).

The exact mechanism linking import perturbation and the neuronal irregularities, including the role of TNTs and rescue, remains unclear. Nevertheless, the evidence suggests that our artificial trapping substrate is indeed mimicking what is seen in disease. Therefore, it provides us with a useful tool to better understand import dysfunction as a potential contributory factor in the onset of neurodegenerative disease. Moreover, deeper analysis of the potential TNT-mediated rescue mechanism will be interesting with regards to early and late onset neurodegeneration, and perhaps how these pathways could be manipulated to delay the onset of disease.

## Acknowledgments

We gratefully acknowledge access and support of the Wolfson Bioimaging Facility for imaging, and the University of Bristol FACS facility for seahorse analysis. Many thanks to Drs Stephen Cross and Richard Seager for image analysis support and advice. We acknowledge the University of Bristol Animal Services Unit for animal support.

## Author contribution

HIN, JMH and IC designed experiments; HIN conducted all experiments; KAW generated Tau constructs; HIN, JMH and IC wrote the manuscript; IC and JMH secured funding and led the project.

## Declaration

The authors declare no competing interests.

## Data and materials availability

All data are available in the main text or the supplementary materials.

## Supplementary Information

## Materials & Methods

### Reagents

All chemicals were of the highest grade of purity commercially available and generally purchased from Sigma, UK unless stated otherwise. Aqueous solutions were prepared in ultrapure water, while for non-aqueous solutions, ethanol or DMSO was used as solvent instead. Antibody catalogue numbers and suppliers are detailed in figure legends.

### Generation of Constructs

Constructs were generated by standard cloning techniques. Briefly, PCR reactions were carried out using Q5 High Fidelity Hot Start DNA Polymerase (New England Biolabs (NEB)), using 20 pmol primers and 200 pg template DNA, as per manufacturers’ instructions. PCR products were purified using QIAquick PCR Purification Kit (QIAgen). Restriction digest reactions were carried out using NEB restriction enzymes at 37°C for 45-60 min. Ligation reactions were carried out using T4 DNA Ligase (NEB) overnight at 16°C. Transformation was carried out in *E. coli* cells (α-select, XL1-Blue, or BL21-DE3 cells were used, depending on application; all originally sourced from NEB) for 30 min on ice, followed by heat shocking (45 sec, 42°C), and a further 15 min on ice. Cells were recovered by incubating in LB media at 37°C for 1 h and then plated on LB-agar plates containing appropriate antibiotic. Plasmids were prepared by mini or maxi preps using commercially available kits (Qiagen and Promega, respectively) following manufacturers’ instructions, and verified by DNA sequencing using Eurofins Genomics TubeSeq service.

Specifically, human Tau 4R0N isoform WT or P301L coding sequences were amplified from pSinRep5-EGFP-Tau expressing plasmids (a gift from Prof Neil Marrion, University of Bristol) by PCR and cloned with an N-terminal Myc-tag into the SpeI and BamHI sites of the lentiviral plasmid pXLG3-PX-WPRE.

### Protein Purification

Protein expression was carried out as described previously (Pereira et al., 2019). A single colony of transformed BL21 (DE3) bacteria was grown in LB with appropriate antibiotic overnight (37°C; 200 rpm). Pre-cultures were used to inoculate a secondary culture at a 1:100 dilution, in 2X YT supplemented with appropriate antibiotic. Secondary cultures were grown until mid-log phase, then induced with 1 mM IPTG or 0.2% (w/v) arabinose and grown for a further 3 h. Cells were harvested by centrifugation (15 min; 6000 xg), resuspended in TK buffer (20 mM TRIS base, 50 mM KCl; pH 8.0), cracked in a cell disruptor (Constant Systems; 2 cycles at 25 kpsi), and clarified by centrifugation (45 min; 38,000 rpm).

#### GST-tagged Recombinant Perfringolysin (rPFO)

Supernatant (soluble fraction) was loaded onto a 5 ml GSTrap 4B column (GE Healthcare) and the column was washed with TK buffer until the absorbance of the flow through ceased to decrease any further. The peptide was eluted using 10 μM reduced glutathione, prepared fresh. Eluted fractions were pooled and loaded onto a 5 ml anionic exchanger (Q- column; GE Healthcare). A salt gradient of 0-1 M was applied over 20 min and the protein was eluted in 5 ml fractions. The fractions containing the protein were confirmed by SDS-PAGE with Coomassie staining, then pooled and spin concentrated. The final protein concentration was determined based on an extinction coefficient of 117,120 M^−1^ cm^−1^. The protein was aliquoted, snap frozen, and stored at −80°C until required. For each assay, a fresh aliquot was thawed, used immediately, and discarded afterwards, due to the instability of this protein (Lei & Bochner, 2021).

#### GST-Dark Peptide

GST-Dark was prepared as described previously (Pereira et al., 2019). Supernatant was loaded onto a GSTrap 4B column and purification was carried out exactly as for rPFO but without the ion exchange chromatography. Analysis, yield, and freezing was carried out exactly as for rPFO. Protein concentration was determined based on an extinction coefficient of 48,360 M^−1^ cm^−1^.

#### His-tagged Su9-EGFP-pep86

Inclusion bodies (insoluble fraction) were solubilised in TK buffer supplemented with 6 M urea, before loading onto a 5 ml HisTrap HP column (GE Healthcare). The protein was eluted in 300 mM imidazole. Eluted fractions containing the desired protein were pooled and loaded onto a 5 ml cationic exchanger (S-column; GE Healthcare). A salt gradient of 0-1 M was applied over 20 min and the protein was eluted in 5 ml fractions. Imidazole was removed by spin concentration, followed by dilution in TK buffer containing 6 M urea. Analysis, yield, and freezing were carried out exactly as for rPFO. Protein concentration was determined based on an extinction coefficient of 28,880 M^−1^ cm^−1^.

### Cell Culture

#### HeLa/HEK293T Cell Culture

HEK293T (ECACC) and glucose-grown HeLa cells (HeLaGLU; ATCC) were maintained in Dulbecco’s Modified Eagle’s Medium (DMEM; Gibco; 41965039) supplemented with 10% (v/v) foetal bovine calf serum (FBS; Invitrogen) and 1% (v/v) penicillin-streptomycin (P/S; Invitrogen). When OXPHOS dependence was required, HeLa cells were cultured in galactose medium (HeLaGAL; ATCC) consisting of DMEM without glucose (Gibco; 11966025) supplemented with 10 mM galactose, 1 mM sodium pyruvate, 10% FBS and 1% P/S. Cells were cultured in galactose media for at least 3 weeks prior to experiments on HeLaGAL cells. Cells were maintained in T75 ventilated flasks in humidified incubators at 37°C with 5% CO_2_.

#### Primary Neuronal Cell Culture

Primary neuronal culture was carried out following established lab protocols, as described previously (Martin & Henley, 2004). Briefly, primary hippocampal and cortical neurons were isolated from embryonic day 18 (E18) Han Wistar rat pups. Pregnant Han Wistar rats, bred in-house at the University of Bristol Animal Services Unit, were anaesthetised using isoflurane with pure oxygen flow and humanely killed by means of cervical dislocation, following Home Office Schedule 1 regulations. Isolated cortices and hippocampi were washed extensively in HBSS and dissociated by incubation with 10% (v/v) Trypsin-EDTA solution at 37°C for 15 and 9 min, respectively. Cells were grown in plating medium (Neurobasal medium (Gibco) supplemented with 5% (v/v) horse serum (Sigma), 2% (v/v) B27 (Gibco), 1% P/S, and 5 mM Glutamax (Gibco)). After 3 h, plating medium was removed and replaced with feeding medium (Neurobasal medium supplemented with B27, P/S, and 2 mM Glutamax). For biochemistry, cortical neurons were seeded at a density of 500,000 cells per well and for imaging experiments, hippocampal neurons were seeded at a density of 150,000 cells per well, where one well represents a ~35 mm surface. All plates (containing sterile coverslips if appropriate) were incubated with poly-L-lysine (0.5 mg/ml or 1 mg/ml for plastic or glass, respectively, in sterile borate buffer (1 mM borax and 5 mM boric acid)) overnight, extensively washed with tissue culture grade H_2_O and incubated overnight in plating medium prior to plating cells, to aid adhesion. Cells were maintained in humidified incubators at 37°C with 5% CO_2_.

### Transfection of Cells

#### HeLa Cell Transfection

HeLa cells were plated and grown up to ~70-80% confluency. At this point cells were transfected with 1 μg (per 35mm dish) of the desired DNA using Lipofectamine 3000 reagent (Thermo Fisher Scientific), at a 1:1.5 ratio of DNA:Lipofectamine, following the manufacturers’ protocol. Cells were then grown for a further 24-72 h prior to experimental analysis.

#### Primary Neuron Transfection

Primary neurons were transfected at 14 days *in vitro* (DIV14) with 0.5-1 μg of the desired DNA using Lipofectamine 2000 reagent (Thermo), at a ratio of 1.5 μl Lipofectamine 2000 per 1 μg DNA. Coverslips with adhered neurons were washed in plain neurobasal medium and transferred to dishes containing 1 ml plain neurobasal medium, to which the transfection mix was added. 45 min after transfection, coverslips were washed again to remove all remaining DNA and Lipofectamine 2000 from cells and returned to their conditioned growth medium. Neurons were then maintained for a further 7 days prior to fixation.

### Cell Transduction by Lentiviral Infection

Lentiviral particles were produced in HEK293T cells by addition of a mixture of DNAs (27.2 μg DNA to be produced, and packaging vectors pMDG2 (6.8 μg) and pAX2 (20.4 μg)) and pEI transfection reagent reagent (1.5:1 pEI:DNA to be produced) in OptiMEM medium (Gibco) to HEK293T cells in a T75 flask, followed by incubation for 6 hours at 37°C, 5% CO_2_. Media was then changed to complete DMEM, and cells incubated for 72 h to allow lentivirus particle production. Lentivirus particles were harvested at 48 and 72 h for maximum yield, pooled, spun down at 4000 xg for 5 min to remove dead cells, and concentrated by adding Lenti-X concentrator (Takara Bio) at a 1:3 ratio and incubating at 4°C for at least 1 h. Lentivirus was then pelleted by centrifugation at 4000 xg for 45 min. Pellets were resuspended in plain DMEM at 1:50 of initial supernatant volume and aliquoted and stored at −80°C until required. For infection, concentrated lentivirus (volume optimised by titration of each fresh batch) was added dropwise to cell media and cells incubated prior to experimental analysis.

### Mitochondrial Isolation

Confluent cells were harvested by trypsinisation, pelleted, and washed extensively with HBSS. Pellets were frozen overnight at −80°C and thawed the next day, to weaken membranes. Subsequently, mitochondrial isolation was performed using Mitochondrial Isolation Kit for Cultured Cells (Abcam; ab110170) following manufacturers’ instructions. Mitochondrial protein concentration was calculated by BCA assay (Pierce^TM^ BCA Protein Assay Kit, Thermo), following manufacturers’ instructions, and using BSA as a standard.

### Immunoprecipitation

Following mitochondrial isolation, mitochondria were gently lysed using 4.5 g GDN/ g protein in IP buffer (0.1M TrisHCl, 0.15M NaCl, phospholipids (0.03 mg/ml PE; 0.03 mg/ml PG; 0.09 mg/ml PC), 1X cOmplete ULTRA Protease Inhibitor Cocktail). Myc-tagged proteins were isolated using 10 μl Myc-Trap beads (Chromotek). Supernatant (lysed mitochondrial sample) was incubated on a rotating wheel with beads overnight at 4°C. Subsequently, beads were washed in IP buffer. After washing, supernatant was removed, and samples were analysed by Western blotting.

### Total Protein Cell Lysis

For extraction of total protein lysate, cells were washed extensively with HBSS, and RIPA buffer (Sigma, supplemented with 1 mM PMSF) was added (200 μl for ~35 mm surface, scaled up or down appropriately). Cells were scraped on ice into Eppendorf tubes, which were incubated for 1 h on a rotating wheel at 4°C, prior to centrifugation at 10,000 xg for 15 min at 4°C. Protein samples were stored at −20°C. Protein concentration was determined by BCA assay.

### Western Blotting

Following protein extraction, samples were heated to 95°C for 5 min in the presence of LDS supplemented with 50 mM DTT. 30 μg of total protein was loaded on 4-12% BOLT gels (Thermo), separated (200 V, 24 min), and transferred onto polyvinylidene difluoride (PVDF) membranes (activated with methanol; Thermo) with transfer buffer (336 mM tris, 260 mM glycine, 140 mM tricine, 2.5 mM EDTA) using a semi-dry Pierce Power Station transfer system (Thermo; 2.5 mAmp, 10 min). Membranes were blocked for 1 h in milk (5% w/v in TBS-T: 20 mM TRIS, 1.5 M NaCl, 0.1% (v/v) Tween-20 (pH 7.6)) and incubated in 5% milk containing the appropriate primary antibody (4°C, overnight; see figure legend). Membranes were washed extensively with TBS-T and probed with appropriate secondary antibody in 2.5% milk (RT, 1 hour). Membranes were washed with TBS-T, incubated with ECL substrate (GE Healthcare), and developed using an Odyssey Fc Imaging System (LI-COR). Analysis and quantification were carried out using Image Studio Lite software.

### NanoLuc Assay

NanoLuc assays were carried out exactly as described previously (Hope I. Needs et al., 2022). Briefly, cells expressing *eqFP670-P2A-Cox8a-11S* and plated on standard white flat-bottom 96-well plates were washed with HBSS and incubated in HBSS wash buffer (HBSS supplemented with 5 mM D-(+)-glucose, 10 mM HEPES (Santa Cruz Biotechnology, Germany), 1 mM MgCl_2_, 1 mM CaCl_2_; pH 7.4). A fluorescence read was taken at 605/670 nm using monochromators with gain set to allow maximum sensitivity without saturation, using a BioTek Synergy Neo2 plate reader. Cells were transferred to permeabilised cell assay master mix (225 mM mannitol, 10 mM HEPES, 2.5 mM MgCl_2_, 40 mM KCl, 2.5 mM KH_2_PO_4_, 0.5 mM EGTA; pH 7.4) supplemented with 5 mM succinate, 1 μM rotenone, 0.1 mg/ml creatine kinase, 5 mM creatine phosphate, 1 mM ATP, 0.1% (v/v) Prionex, 3 nM rPFO (purified in house), 20 μM GST-Dark (purified in house), 1:800 furimazine (Nano-Glo® Luciferase Assay System; Promega). A baseline read of 30 seconds of background luminescent signal was taken prior to injection of purified substrate protein (*Su9-EGFP-pep86*) to 1 μM final concentration, followed by a further bioluminescence read corresponding to import, lasting 30 min. Bioluminescence was read using a BioTek Synergy Neo2 plate reader (Agilent, UK) or a CLARIOstar Plus plate reader (BMG LabTech, UK) without emission filters with gain set to allow maximum sensitivity without saturation, and with acquisition time of 0.1 sec per well. Row mode was used, and reads were taken every 6 sec or less, with wells in triplicate.

### Confocal Microscopy

#### Fixed Cell Confocal Microscopy

Cells on glass coverslips were washed in HBSS and fixed in 4% (v/v) paraformaldehyde (PFA) with 2% (w/v) sucrose (sucrose used for neurons only, to retain osmolarity), for 12 min at 37°C. Coverslips were then washed extensively in HBSS, and PFA was quenched using 100 mM Glycine. Cells were then stained for ICC or mounted.

For immunocytochemistry, following fixation and quenching, cells were permeabilised by incubation in 0.1% (v/v) Triton-X in PBS for 5 min, washed in PBS and blocked in 3% (w/v) BSA in PBS for 30 min. Coverslips were incubated on a drop of primary antibody in 3% BSA overnight at 4°C. Coverslips were washed extensively in PBS and incubated with the appropriate secondary antibody (2 hours, RT) in 3% BSA, then washed again.

Coverslips were dipped in ddH_2_O and mounted onto slides using Fluoromount-G mounting medium (Thermo). Coverslips were left to dry overnight and imaged using a Leica confocal microscope (SP5II or SP8) and LAS X software platform. Laser lines used were 405, 488, 562, and 633 nm, with gain set to allow maximum sensitivity without saturation, and z-stacks were taken.

#### Live Cell Confocal Microscopy

Transfected cells were grown to confluency on 35 mm glass bottomed dishes (Corning). Cells were washed in HBSS and incubated in HBSS imaging buffer (HBSS supplemented with 5 mM Glucose, 10 mM HEPES, 1 mM MgCl_2_, 1.26 mM CaCl_2_) with 25 nM tetramethylrhodamine (TMRM) for 30 min at 37°C. Cells were imaged immediately, with TMRM retained in the buffer, using a Leica SP8 confocal microscope and LAS X software platform, at 37°C.

### Image Analysis

All image analysis was performed using the FiJi image processing package (Rueden et al., 2017; Schindelin et al., 2012). Macros were written by Dr Stephen Cross (Wolfson Bioimaging Facility, University of Bristol), within the FiJi plugin Modular Image Analysis (MIA) package version 0.21.0, which is publicly accessible on GitHub with a linked version specific DOI from Zenodo (Cross, 2021). Where data was analysed manually, data was blinded prior to manual analysis.

#### Mitochondrial Morphology Analysis

Mitochondrial pre-processing, as well as branch and network analysis was carried out as described previously (Cross & Seager, 2022; Seager, 2020), using a FiJi plugin adapted from the mitochondrial network analysis (MiNA) toolset (Valente et al., 2017). Briefly, for pre-processing, cells of interest were outlined and outside cleared, prior to z-stack max intensity projection, contrast enhancement and background subtraction. Median, unsharp, and tubeness filters were applied. For branch/network analysis, images were binarised and skeletonised, and the skeleton analysed to obtain information on the branches and pixels, allowing analysis of mitochondrial branching as a readout of mitochondrial network complexity and fragmentation.

#### Membrane Potential Analysis

TMRM analysis of mitochondrial membrane potential was done based on a published protocol (Creed & McKenzie, 2019). Briefly, an average of the first 10 frames (TMRM fluorescence where images were captured by live confocal microscopy over 2 minutes) was taken, then the mitochondria were thresholded and their intensity measured. The average of the final three frames (after carbonyl cyanide m-chlorophenylhydrazone (CCCP) treatment to dissipate the ΔΨ) was subtracted as a control and the intensity of the subtracted image was re-measured.

#### Neuronal Morphology Analysis

Analysis of neuronal complexity (number of processes, *i.e.,* axon and dendrites) was carried out by manually counting the number of processes protruding from the soma (at a given distance from the soma – highlighted by circles on images). This was carried out on blinded data, where expression of the protein of interest had been confirmed (prior to data blinding) and all processes were highlighted using an mCherry cytosolic marker.

Initial segments of axons in transfected cells were identified by staining for Ankyrin G. Following identification of the axon to be measured, the length of the axon was measured by tracing the length of the axon, highlighted by the mCherry cytosolic marker, using the Simple Neurite Tracer (SNT) component of the Neuroanatomy plugin in FiJi.

Synapses were counted manually. First, data was blinded, and dendrites were identified. The dendrite of interest was measured using the SNT plugin and the synapses along the region of interest were counted manually using the PSD95 stain as reference. A total of 5 dendrites were analysed per cell and an average was taken for each cell.

### Seahorse Assays: Mitochondrial Stress Test

Cells were seeded in 6-well plates (200,000 cells/well) and infected with the appropriate lentivirus, as described in figure legends. The day prior to the assay, cells were detached by trypsinisation, counted, and seeded at 10,000 cells/well in 96 well Seahorse XF cell culture plates (Agilent). The sensor cartridges were hydrated overnight with tissue culture grade H_2_O in a non-CO_2_ incubator at 37°C as per manufacturer’s instructions. On the day of the assay, H_2_O in the sensor plate was replaced with Seahorse XF Calibrant (Agilent) and cells were washed with HBSS and incubated in Seahorse media (Seahorse XF assay medium (Agilent), 1 mM pyruvate (Agilent), 2 mM glutamine (Agilent), 10 mM D-(+)-galactose). Both sensor and cell plates were incubated in a non-CO_2_ incubator at 37°C for 1 h prior to the assay. The sensor plate was loaded with oligomycin (15 μM for 1.5 μM final concentration in wells; injector A), CCCP (5 μM for 0.5 μM final; injector B), and antimycin A and rotenone (5 μM/5 μM for 0.5 μM/0.5 μM final; injector C). The sensor plate was calibrated prior to loading cells and running a mitochondrial stress test using a Seahorse XFe96 (Agilent). Following assays, cells were washed and fixed in 1% (v/v) acetic acid in methanol at −20°C overnight for SRB assays, which were used for normalising data to protein content.

### SRB Assay

Cells were fixed with ice cold 1% acetic acid in methanol overnight at −20°C. Fixative was aspirated and plates were allowed to dry at 37°C. SRB (0.5% (w/v) in 1% (v/v) acetic acid in dH_2_O) was added to cover cells, and plates incubated for 30 min at 37°C. SRB was then aspirated, and unbound stain removed by washing 4 times with 1% acetic acid in dH2O, prior to drying plates at 37°C. Bound protein stain was solubilised by shaking incubation with 10 mM Tris (pH 10; 15 min, RT). Absorbance was read on a microplate reader with a 544/15 nm filter.

### Statistical Analysis

Statistical significance between groups was determined using unpaired Student’s t-Test or one or two-way ANOVA with interaction if more than two groups were analysed. ANOVA was nested if multiple objects were analysed per biological replicate. Following ANOVA, p values were adjusted for multiple comparisons through Tukey’s *post hoc* test and differences were considered significant at 5% level. Statistical analyses were performed using Graph Pad Prism version 9 (GraphPad Software, Inc., San Diego, CA, USA).

**Fig. S1.**
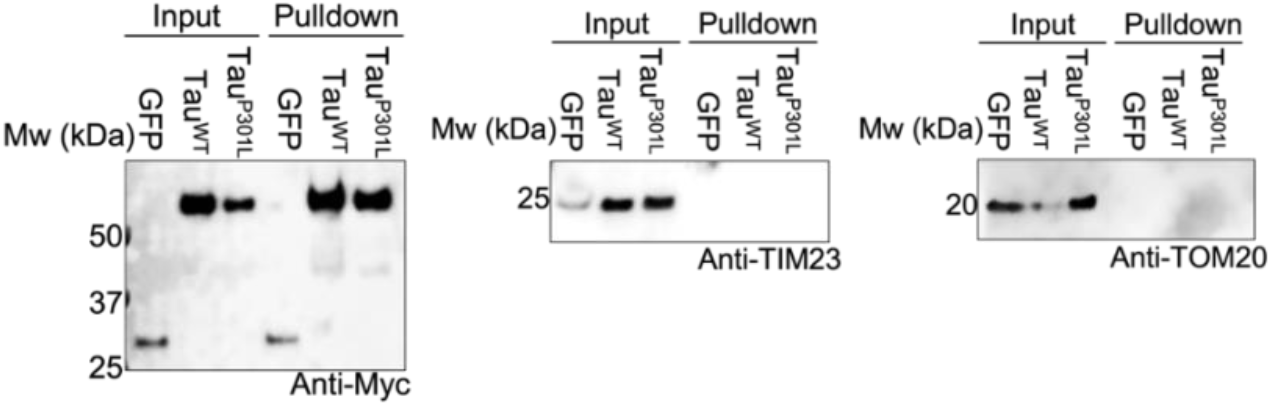
Representative Western blot showing levels of Myc-tagged proteins of interest (showing pulldown worked), as well as TIM23 and TOM20, in input and pulldown samples from the mitochondrial fraction of HeLaGAL cells overexpressing Myc-tagged GFP, Tau^WT^, or Tau^P301L^. Corresponding to IP experiment shown in Fig. 1D. N=4 biological replicates.

**Fig. S2.**
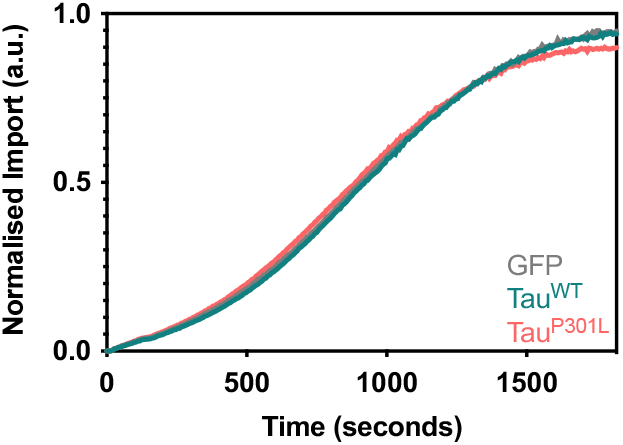
NanoLuc import assay trace showing import of precursor protein *Su9-EGFP-pep86* in HeLaGAL cells expressing *Cox8a-11S* and GFP, Tau^WT^, or Tau^P301L^. Averaged, normalised traces are shown (normalised to eqFP670 expression and max amplitude/run), error bars represent SD. N=3 biological repeats, each with n=3 technical replicates. Corresponds to quantification of import amplitude shown in Fig. 1E.

**Fig. S3.**
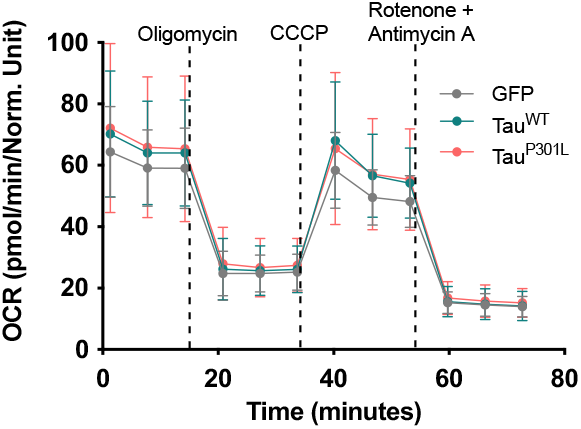
Mitochondrial stress test showing oxygen consumption rate (OCR) of HeLaGAL cells expressing Myc-GFP, Myc-Tau^WT^, or Myc-Tau^P301L^. Data is normalised to protein content as determined by SRB assays. Error bars show SD. N=6 biological replicates, each with n=3 technical replicates.

**Fig. S4.**
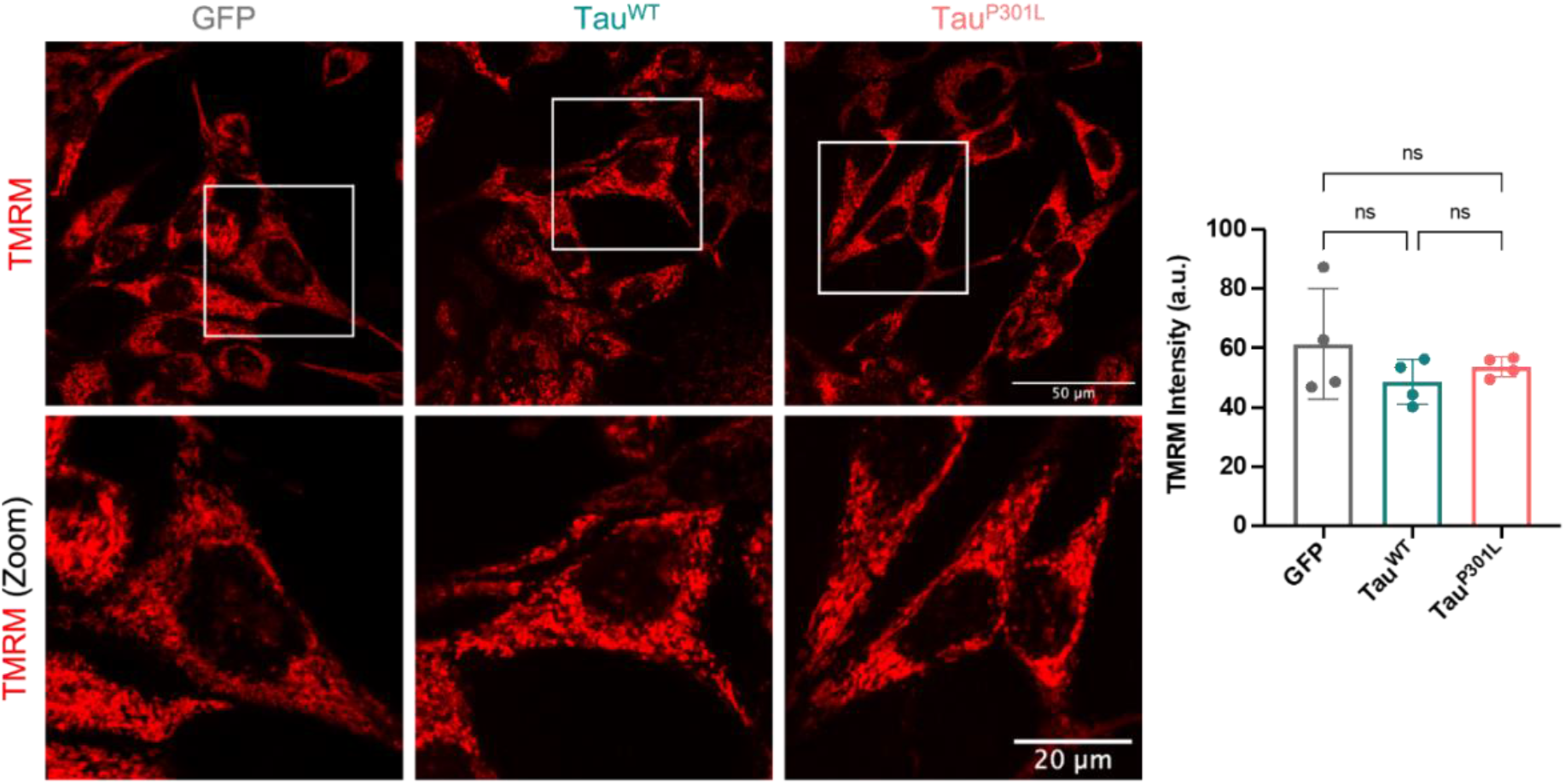
Representative confocal images (left) showing TMRM fluorescence (red) in the mitochondria of HeLaGAL cells expressing Myc-GFP, Myc-Tau^WT^, or Myc-Tau^P301L^. CCCP was added after 2 minutes to control for background fluorescence. TMRM intensity was quantified (right) using a FiJi macro (see Methods). N=4 biological replicates. Error bars show SD. One-way ANOVA and Tukey’s *post hoc* test were used to determine significance.

**Fig. S5.**
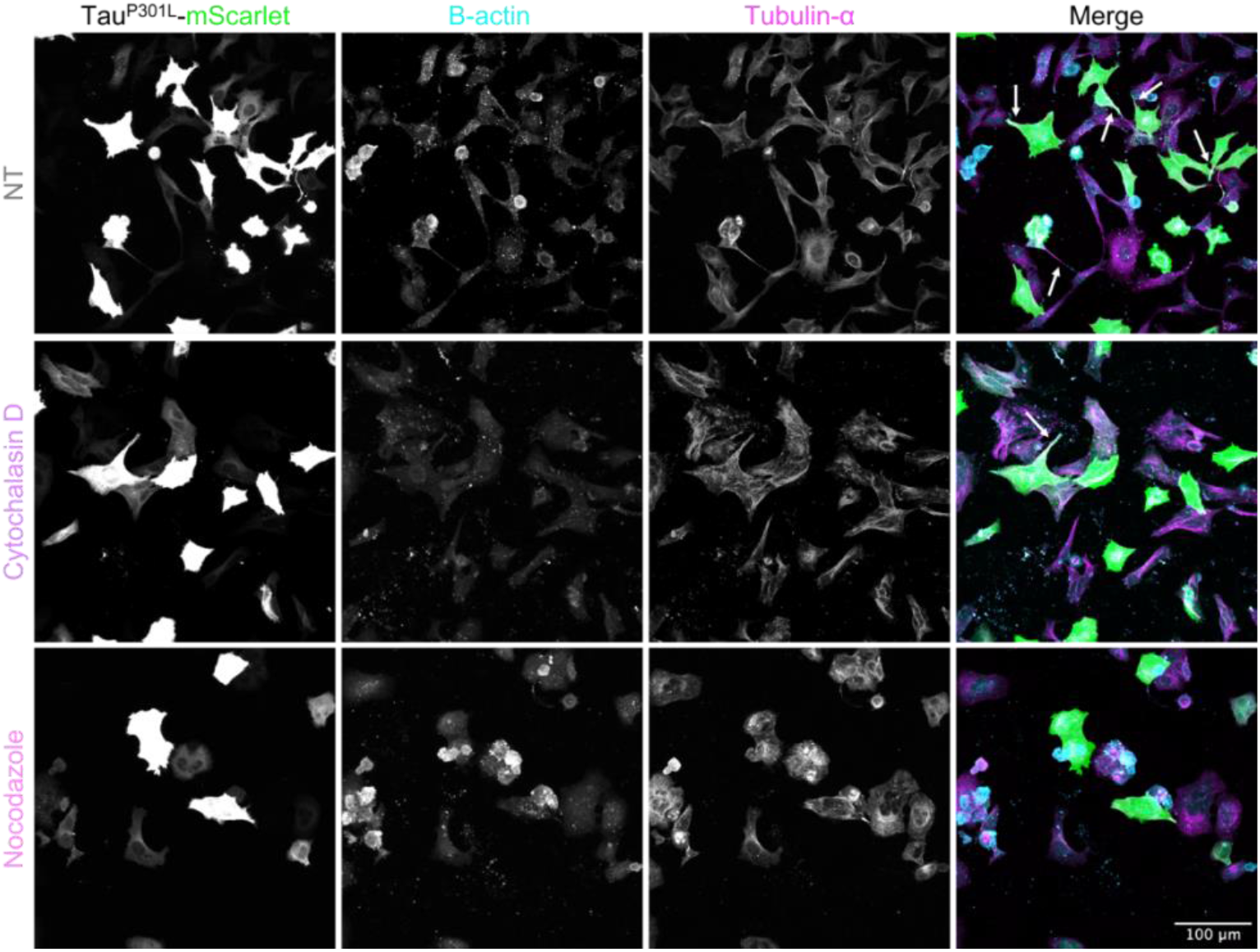
Representative confocal images of HeLaGAL cells exposed to Tau^P301L^ overexpression in the presence of TNT inhibitors. HeLaGAL cells were subjected to overexpression of Tau^P301L^-mScarlet (green) and either untreated (NT; DMSO only) or treated with 50 nM Cytochalasin D or 100 nM Nocodazole (48 h). Cells were fixed and stained for β-actin (cyan) and tubulin-α (magenta; BioRad MCA78G). Arrows in merge highlight TNTs. N=3 biological replicates. Quantification shown in Fig. 2D.

**Fig. S6.**
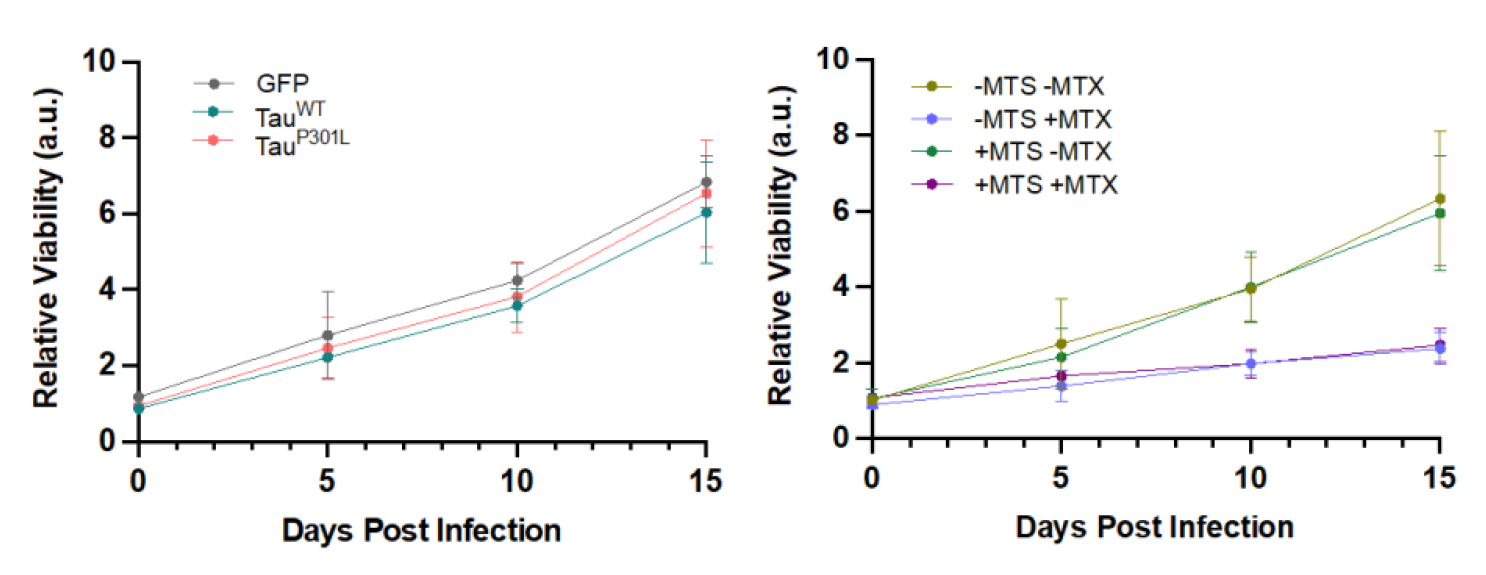
Viability of primary cortical neurons lentivirally expressing GFP (control), Tau^WT^ or Tau^P301L^ (left), or EGFP-DHFR(-MTS) or Su9-EGFP-DHFR (+MTS) +/− MTX (right), as determined by cell density, quantified by SRB assays. Data was normalised to the average reading for the 0 days post infection timepoint, to give relative viability of the neurons in response to the varying expression/treatment. N=4 biological replicates. Error bars show SD.

**Fig. S7.**
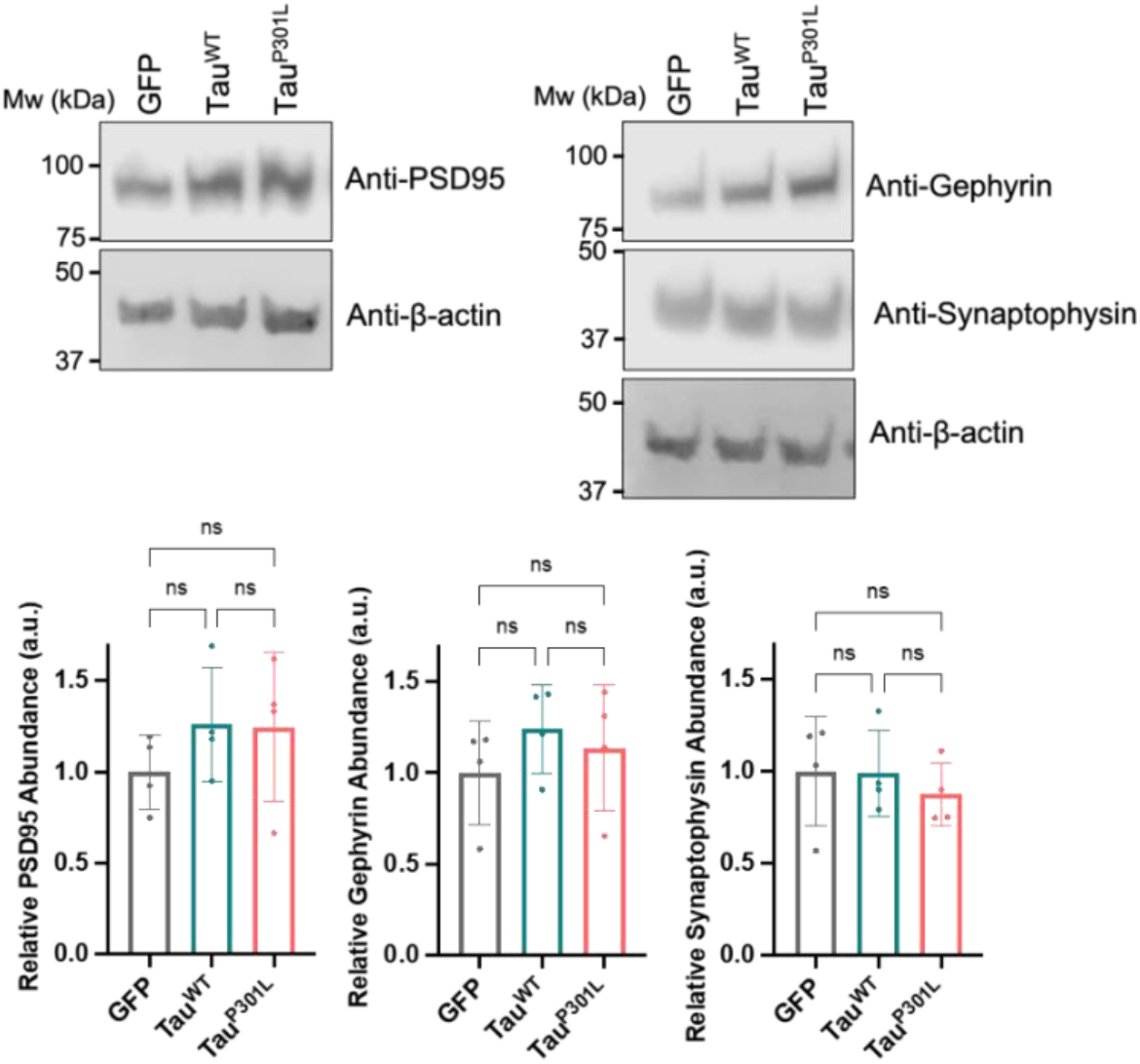
Representative Western blots showing levels of PSD95, Gephyrin (Synaptic Systems; 147111), and Synaptophysin (Merck Millipore; 573822) in DIV21 cortical neurons after 7 days of lentiviral expression of Myc-GFP, Myc-Tau^WT^, or Myc-Tau^P301L^. β-actin was used as a loading control. N=4 biological replicates. Histograms show quantification of PSD95, Gephyrin, and Synaptophysin relative abundance, normalised to β-actin. Error bars show SD. One-way ANOVA and Tukey’s *post hoc* test were used to determine significance.

**Fig. S8.**
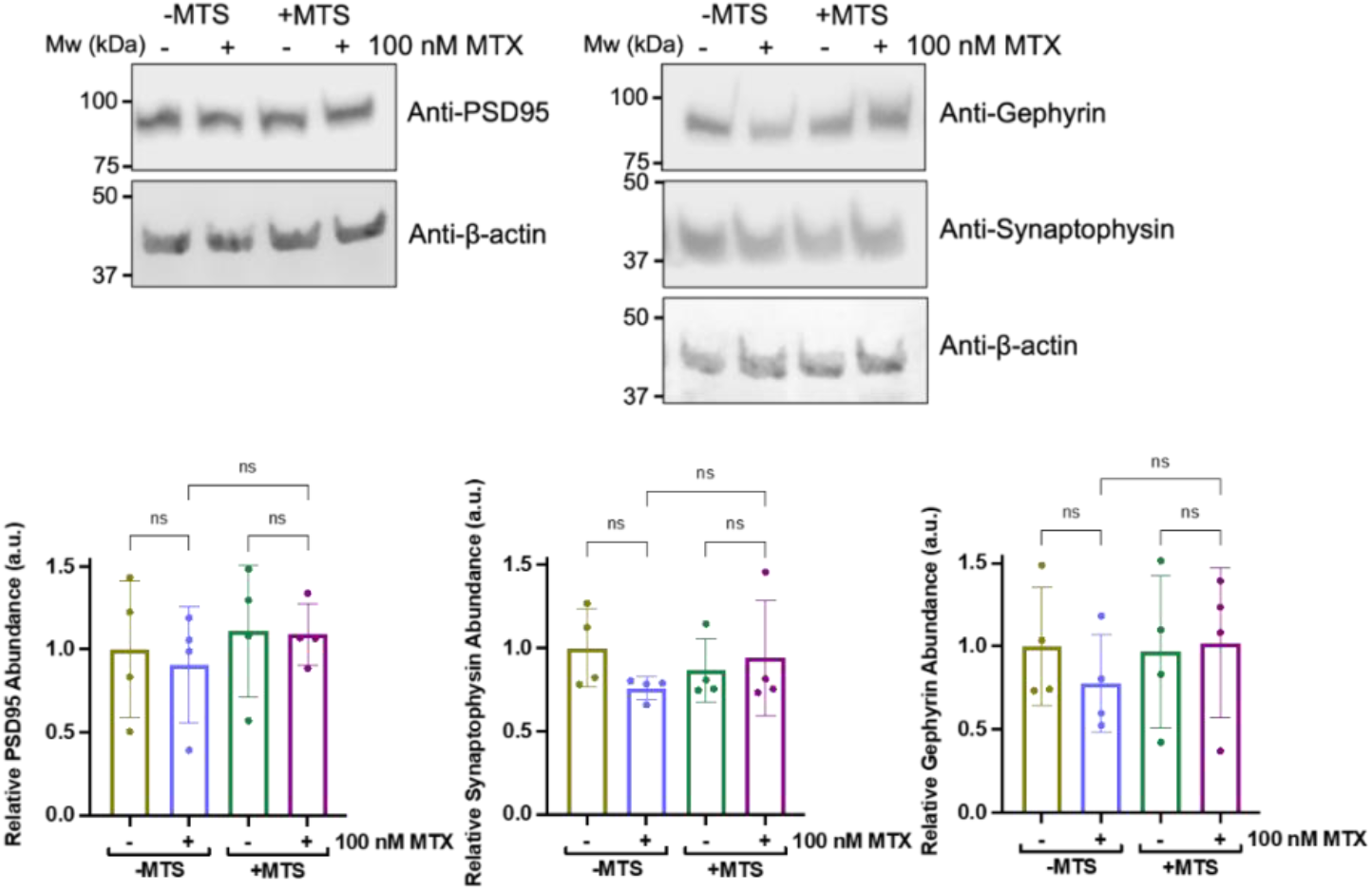
Representative Western blots showing total cellular abundance of PSD95, Gephyrin, and Synaptophysin in DIV21 cortical neurons after 7 days of lentiviral expression of EGFP-DHFR (-MTS) or Su9-EGFP-DHFR (+MTS) +/− MTX. β-actin was used as a loading control. N=4 biological replicates. Histograms show quantification of PSD95, Gephyrin, and Synaptophysin relative abundance, normalised to β-actin. Error bars show SD. One-way ANOVA and Tukey’s *post hoc* test were used to determine significance.

